# Megakaryocytic proteomic signatures define myeloproliferative neoplasms

**DOI:** 10.64898/2026.10.30.762990

**Authors:** Indu Raman, Anne Pettikiriarachchi, Ricardo Ataíde, Surender Juneja, Yu Zhu, Ellen Tsui, Samantha J Emery-Corbin, Sukhdeep Spall, Jumana Yousef, Laura F Gianni, Pradeep Rajasekhar, Allan Motyer, Costas K Yannakou, Sant-Rayn Pasricha, Cavan Bennett

**Affiliations:** Walter and Eliza Hall Institute of Medical Research, Melbourne, Australia; Epworth HealthCare, Melbourne, Australia; The University of Melbourne, Parkville, Victoria, Australia; Royal Melbourne Hospital, Melbourne, Australia; Peter MacCallum Cancer Centre, Melbourne, Australia; Department of Biochemistry and Molecular Biology, Biomedicine Discovery Institute, Monash University, Clayton, Victoria, Australia

## Abstract

The classical Philadelphia-negative myeloproliferative neoplasms (MPNs) are clonal stem cell disorders of the bone marrow with significant niche dysregulation driven in part by abnormal megakaryocytes. The three classical MPNs: polycythemia vera (PV), essential thrombocythemia (ET) and primary myelofibrosis (PMF), display established megakaryocytic morphological differences which aid in their diagnostic classification. The paucity of megakaryocytes in the bone marrow aspirate has previously limited a proteomic analysis of megakaryocytes in clinical samples. We aimed to undertake an unbiased proteomic analysis of *in situ* megakaryocytes from the bone marrow of patients with PV, ET, and PMF. Using a novel, deep visual proteomics strategy, we dissected and analyzed megakaryocytes from 60 MPN patients and 20 controls (patients with limited stage non-Hodgkin lymphoma who had uninvolved bone marrow). We reveal that megakaryocyte proteomics from MPN patients are distinct to those from control patients, and further, that the megakaryocytes from the three MPNs have unique, disease-specific signatures. The disease-specific megakaryocyte proteome highlights novel pathways of disease.

**Key Points:**

- A novel method can evaluate the proteome of megakaryocytes *in situ* from archived clinical FFPE bone marrow trephine samples
- Megakaryocytes in myeloproliferative neoplasms display distinctive disease-specific functional proteomic signatures

## Introduction

The classical Philadelphia-negative myeloproliferative neoplasms (MPNs) are a related but distinct group of chronic myeloid malignancies comprising three entities: polycythemia vera (PV), essential thrombocythemia (ET), and primary myelofibrosis (PMF). MPNs are driven by constitutively activated JAK-STAT signaling in clonal hematopoietic cells with driver mutations in *JAK2*, *CALR* and *MPL* genes in the majority of cases.^1^

Diagnostic classification into PV, PMF or ET is based on a combination of histological assessment of the bone marrow trephine, including evaluation of megakaryocyte morphology, peripheral blood cell counts, driver mutation status and clinical features.^2^ While MPNs are related disorders it is imperative to distinguish between them since they are fundamentally different disease states with distinct treatment and prognostic implications.^3^ Megakaryocytes are large polypoid hematopoietic cells of the bone marrow that produce platelets, and their distinct spatial and morphological characteristics across the three MPN subtypes form an important part of the diagnostic criteria for each disease. In PV, the megakaryocytes are pleomorphic and lack atypia, while in ET they are numerous and large with hyperlobulated, staghorn-like nuclei.^3^ In PMF they are often densely clustered and demonstrate marked atypia.^3^ Despite these morphologic distinctions, disease classification of MPNs can be challenging, particularly where clinical, hematological and histological features are misaligned.^4,5^ Here, the detectable proteomic differences between megakaryocytes of the three MPN subtypes may be able to simplify these diagnostic challenges in a cost-effective manner.

Clonal megakaryocytes are key drivers of niche dysregulation and disease progression in the MPNs,^6,7^ however the mechanisms underlying these effects remain poorly understood. Proteomic profiling of megakaryocytes in MPNs could present a useful strategy to better understand their pathobiological role and to eventually complement current diagnostic methods. Due to their scarcity in healthy bone marrows (less than 0.05% of total cellularity)^8^ and often hemodiluted or acellular aspirates in MPNs, previous molecular studies have largely relied on megakaryocytes cultured *in vitro* from peripheral blood stem cell collections.^9–11^ Transcriptional and proteomic expression is strongly influenced by the native microenvironment^12–14^, which is highly dysregulated, complex, and incompletely understood in MPNs^7^, making *in vitro* replication of these conditions imprecise and potentially unrepresentative of the *in situ* clinical state.

To characterize the functional proteomic changes within *in situ* megakaryocytes in MPNs, we developed and applied a novel low-input proteomics workflow on archival bone marrow trephine samples of MPN patients with the intention of identifying novel diagnostic and therapeutic biomarkers. We show for the first time that *in situ* differentiated MPN megakaryocytes have an altered proteome compared to control megakaryocytes, and further, that the megakaryocytic proteomes of the three MPNs reflect clearly distinct disease states.

## Methods

Full details are available in the Supplemental Materials.

### Clinical samples and Ethics

Archived formalin fixed paraffin embedded (FFPE) bone marrow trephines obtained at the Royal Melbourne Hospital, Melbourne, Australia between 2009 and 2016 were collected following review and approval of sample collection and experimental methodology by the Melbourne Human Research Ethics Committee (HREC/71841/MH-2020). Cases were included in the study if they met 2022 World Health Organization (WHO) classification criteria for diagnosis of PV, ET and PMF^3^ following a contemporary review of clinical characteristics, peripheral blood counts, morphological assessments of peripheral blood smear, bone marrow aspirate and trephine morphology, and driver mutation status where available, by at least two hematopathologists. Age and sex-matched patients with limited stage B cell lymphoma who had uninvolved bone marrow biopsies collected during the same period were included in the study as controls.

### Deep learning-assisted laser capture microdissection (LCM)-mass-spectrometry based proteomics of in situ megakaryocytes

FFPE bone marrow trephine biopsies were sectioned at 4 µm and mounted onto polyethylene naphthalate (PEN) membrane glass slides (4 µm; Leica, 11600288). CD61 immunohistochemistry was performed following antigen retrieval. Deep learning-based image analysis was carried out using the pixel classifier module of Aivia software (Leica Microsystems, Wetzlar, Germany) to perform automated segmentation, identification and outlining of CD61 stained megakaryocytes. These shapes were subsequently transferred to Leica LMD7 software (version 8.3). Identified shapes were then accurately superimposed upon CD61 positive megakaryocytes on the PEN membrane slide. CD61 positive megakaryocytes were micro-dissected into 96 well Eppendorf plates pre-filled with 30 µL acetonitrile using the Leica LMD 7 system (Leica LMD7 software version 8.3). Following sample preparation, proteins were digested into peptides which were acidified, purified, lyophilized, and reconstituted for LC-MS analysis. Approximately 100 ng peptide was separated by reverse-phase chromatography using a 25-min gradient and analyzed on an Orbitrap Astral mass spectrometer (Thermo) connected to a Neo Vanquish liquid chromatography system (Thermo). Proteomic data were acquired using data-independent acquisition (DIA) with high-resolution MS1 scans and HCD fragmentation. Complete details are given in the Supplemental Materials.

Raw DIA data were analyzed on Spectronaut version 2.0.0 using BGS factory settings of direct DIA analysis. Homo sapiens Uniprot reference proteome (2024) was used for database searching. Data processing and analysis were performed in R (version 4.4.2). Peptide intensities were log_₂_-transformed and summarized to the protein level using limpa (v1.1.6), resulting in a total of 6,476 proteins. RUVIIIC was used for normalization of both datasets. Multivariate analysis using principal component analysis (PCA) was employed to assess overall data structure and identify potential outliers. Differential expression analysis was performed using limma (v3.62.2). Proteins were considered significantly differentially expressed if the false discovery rate (FDR), controlled using the Benjamini–Hochberg procedure, was ≤ 5%. Pathway enrichment analysis using Gene Ontology (GO)^15^, KEGG^16^, and Reactome^17^ were performed on the set of significant proteins.

To identify proteins that discriminate between MPN subgroups, sparse partial least squares discriminant analysis (sPLS-DA) was performed using the mixOmics R package (version 6.32.0)^18^ on normalized levels of 6308 proteins. To determine the optimal number of components and the number of proteins with non-zero loadings in each component, we performed 5-fold cross-validation (CV) with 10 replications.

### Immunohistochemistry and multiplexed immunofluorescence

IHC staining was performed with primary antibody against CDKN1a (p21; rabbit monoclonal clone 12D1, 2947, CST) on the Dako Omnis (Agilent) automated staining platform. OPAL multiplex staining was performed using a BOND RX Fully Automated Research Stainer (Leica Biosystems) using anti-CD61 [Dako M0753], anti-CD41 [Abcam ab134131], anti-PAI-1 [Abcam ab318286] and anti-FGF2 [Abcam ab92337] antibodies. Nuclear staining for CDKN1A and FGF2 in megakaryocytes was assessed by a hematopathologist, who counted at least 500 megakaryocytes in PMF samples and all megakaryocytes in control samples. Staining was quantified as a percentage of megakaryocytes assessed. Trained pixel classifiers in QuPath (version 0.5.1) were used to identify CD41/CD61-positive megakaryocytes and quantify their co-expressed area. A separate classifier distinguished PAI-1-positive cytoplasm, PAI-1-negative cytoplasm, and nuclei within megakaryocytes. This classifier was applied to CD41/CD61 co-expressing areas, and PAI-1 expression was quantified as the percentage of the total CD41/CD61-positive area occupied by PAI-1-positive cytoplasm in each sample. Statistical analysis between control and PMF megakaryocytes was performed using Mann-Whitney U test in Prism Version 10.6.1 (GraphPad Software).

### Data sharing statement

Mass spectrometry proteomics data have been deposited to the Proteome Xchange Consortium via the PRIDE partner repository with the dataset identifier PXD073839. For all other original data contact the corresponding author.

## Results

### A novel workflow for low input proteomics of in situ megakaryocytes in clinical trephine samples

To study the proteomic composition of clinical *in situ* megakaryocytes, we developed a novel workflow for low input proteomics by combining immunohistochemistry, high resolution imaging, deep learning-based image analysis, laser capture microdissection (LCM) and high sensitivity mass spectrometry (Figure 1A). Firstly, 4 µm sections of archival FFPE bone marrow trephines were incubated with anti-CD61 antibodies that were subsequently DAB stained to identify the presence of megakaryocytes (Figure 1B). Approximately 200 CD61+ megakaryocytes per sample were isolated by LCM. Extracted megakaryocyte samples underwent high resolution DIA mass-spectrometry based proteomic analysis.

**Figure 1.**
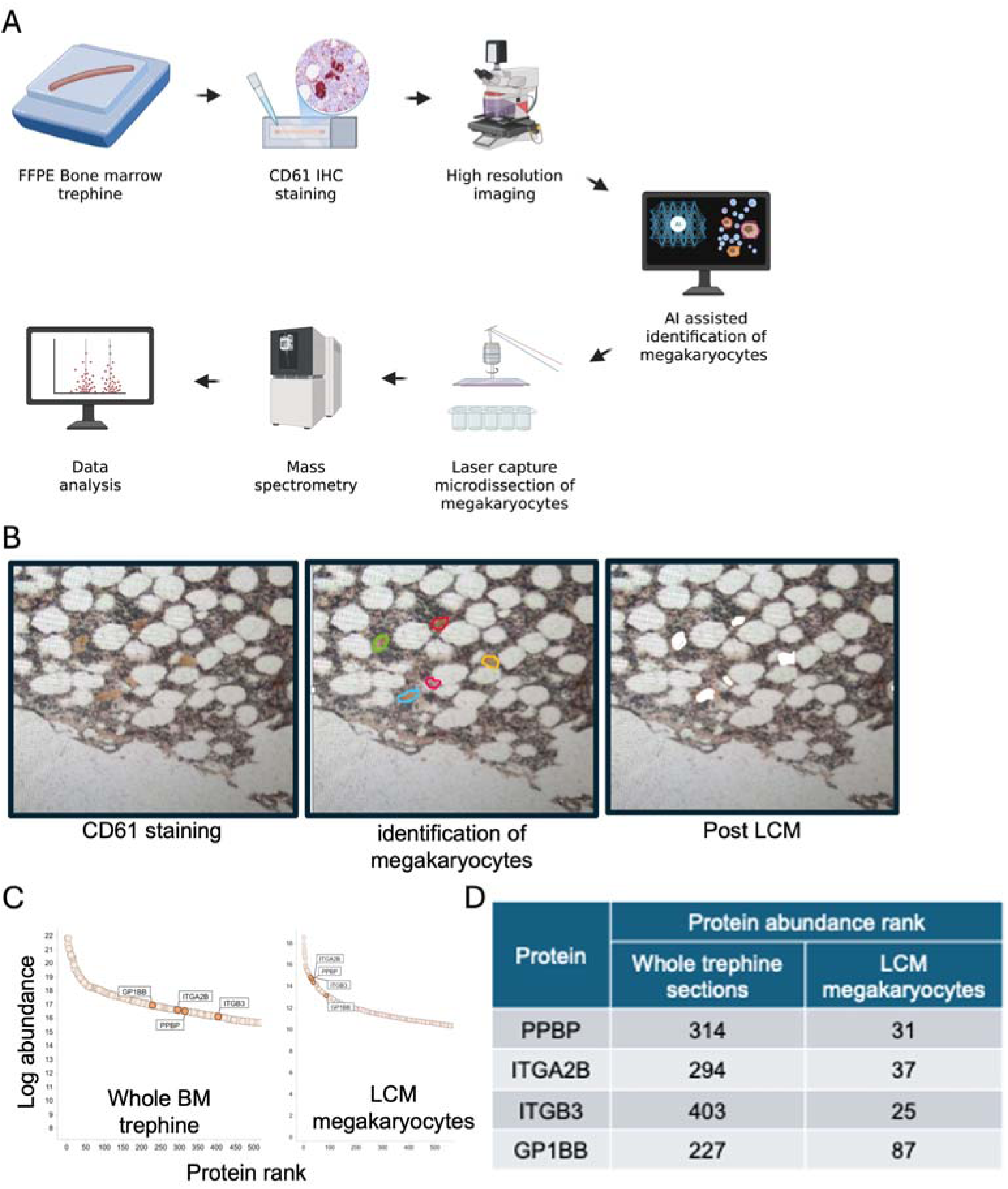
A novel low input proteomics workflow to study *in situ* differentiated megakaryocytes. (A) Methodological overview. (B) CD61 immunohistochemistry of bone marrow trephine (left panel) with deep learning generated shapes around identified CD61+ megakaryocytes (middle panel) and the same section following megakaryocyte laser-capture microdissection (right panel). Log-rank abundance plots of megakaryocyte specific proteins extracted from whole bone marrow section (Left panel) and laser-capture microdissected megakaryocytes (right panel) of control samples. (D) Protein rank of megakaryocyte specific proteins by order of abundance from whole bone marrow section and laser-capture microdissected megakaryocytes proteomic datasets of control samples.

We applied this method to archival bone marrow trephine samples from a cohort of 20 patients with limited-stage non-Hodgkin’s lymphoma who had histologically normal bone marrow and served as a control cohort (Table 1). A total of 6476 proteins were identified. To confirm enrichment of megakaryocytes, we generated a proteomic dataset of whole bone marrow trephine sections from control patient trephines (Supplemental Table 1) and compared the log rank abundance of identified proteins in the LCM-megakaryocyte dataset to that of the whole section dataset. The megakaryocyte-specific proteins, PPBP (CXCL7), ITAG2B (CD41), ITGB3 (CD61) and GP1BB (CD42c), ranked among the top 100 proteins by intensity in the LCM-megakaryocyte dataset but fell between rank 200–500 in the whole-section bone marrow trephine dataset (Figure 1C-D), thereby confirming that our LCM protocol had specifically enriched megakaryocytes.

**Table 1.**
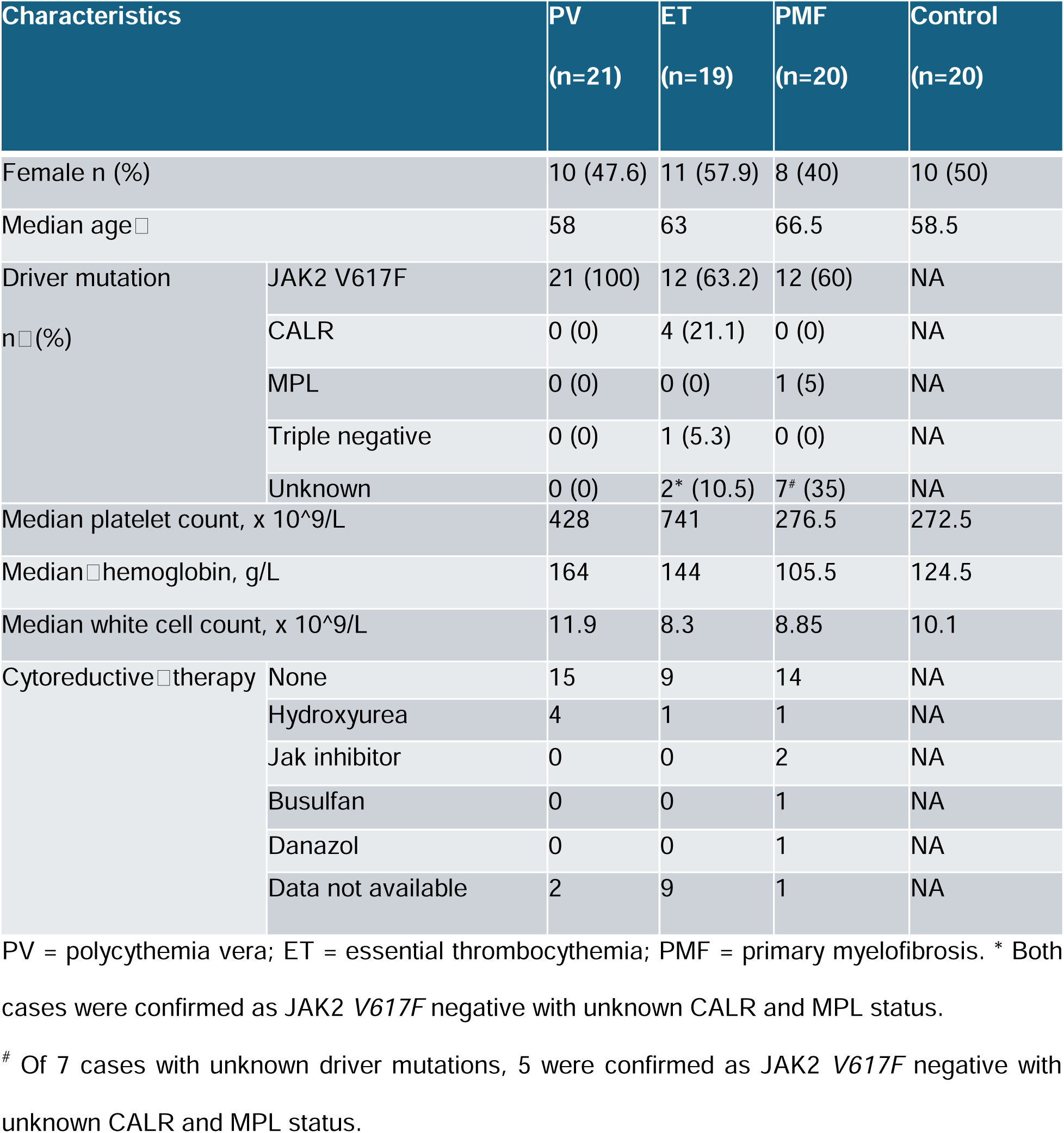
Patient sample characteristics.

| Characteristics |  | PV<br>(n=21) | ET<br>(n=19) | PMF<br>(n=20) | Control<br>(n=20) |
|---|---|---|---|---|---|
| Female n (%) |  | 10 (47.6) | 11 (57.9) | 8 (40) | 10 (50) |
| Median age□ |  | 58 | 63 | 66.5 | 58.5 |
| Driver mutation<br>n□(%) | JAK2 V617F | 21 (100) | 12 (63.2) | 12 (60) | NA |
|  | CALR | 0 (0) | 4 (21.1) | 0 (0) | NA |
|  | MPL | 0 (0) | 0 (0) | 1 (5) | NA |
|  | Triple negative | 0 (0) | 1 (5.3) | 0 (0) | NA |
|  | Unknown | 0 (0) | 2<sup>*</sup> (10.5) | 7<sup>#</sup> (35) | NA |
| Median platelet count, x 10^9/L |  | 428 | 741 | 276.5 | 272.5 |
| Median□hemoglobin, g/L |  | 164 | 144 | 105.5 | 124.5 |
| Median white cell count, x 10^9/L |  | 11.9 | 8.3 | 8.85 | 10.1 |
| Cytoreductive□therapy | None | 15 | 9 | 14 | NA |
|  | Hydroxyurea | 4 | 1 | 1 | NA |
|  | Jak inhibitor | 0 | 0 | 2 | NA |
|  | Busulfan | 0 | 0 | 1 | NA |
|  | Danazol | 0 | 0 | 1 | NA |
|  | Data not available | 2 | 9 | 1 | NA |

PV = polycythemia vera; ET = essential thrombocythemia; PMF = primary myelofibrosis. <sup>\*</sup> Both cases were confirmed as JAK2 *V617F* negative with unknown CALR and MPL status.
<sup>#</sup> Of 7 cases with unknown driver mutations, 5 were confirmed as JAK2 *V617F* negative with unknown CALR and MPL status.

### MPN megakaryocytes have aberrant proteomic signatures

Megakaryocyte morphology is abnormal in MPNs^2^. To understand whether megakaryocytes from the classical MPNs have distinct proteomic profiles, we applied our low input LCM proteomic method to archival bone marrow trephine samples from 60 patients with World Health Organization (WHO)–defined^2^ MPNs - 21 with PV, 19 with ET and 20 with PMF - and controls (Table 1).

We first compared protein expression levels of megakaryocytes from MPN patients to control megakaryocytes. Uniform Manifold Approximation and Projection (UMAP) analysis revealed that MPN megakaryocytes were generally distinct from controls, and strikingly distinct clustering of megakaryocyte proteomes, according to disease subgroup (Figure 2A).

**Figure 2.**
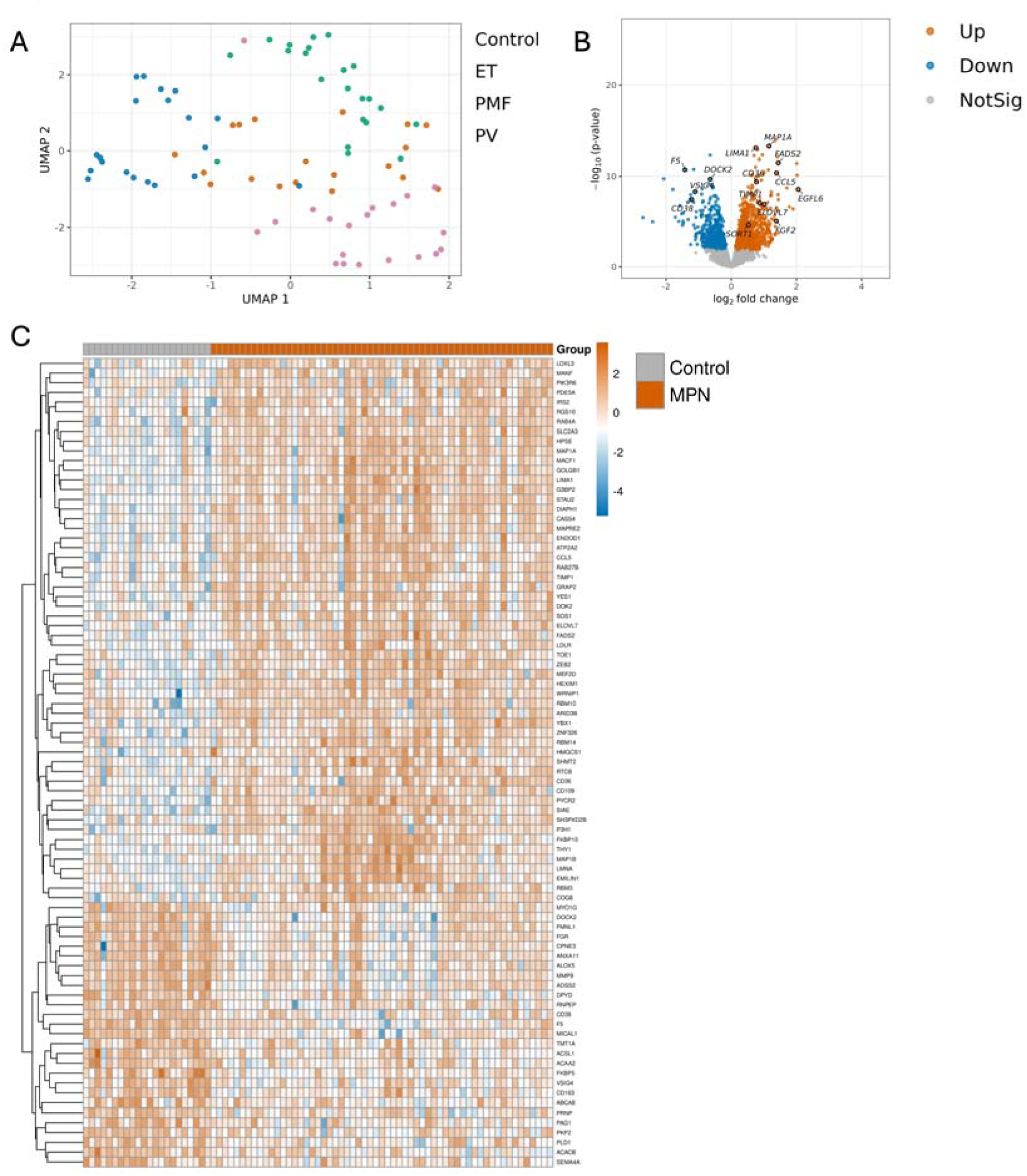
MPN megakaryocytes are proteomically distinct from control megakaryocytes. (A) Uniform Manifold Approximation and Projection (UMAP) projection reveals disease specific clustering. While visual clustering showed overlapping distributions across groups (average silhouette width = 0.031), PERMANOVA confirmed a statistically significant effect of diagnostic group (Control, PV, ET, PMF) on overall proteomic variance (R2 = 0.176, p = 0.001, 999 permutations). (B) Volcano plot visualizing protein expression in MPN and control megakaryocytes. Orange = increased in MPNs; blue = decreased in MPNs; grey = not significantly different. (C) Heatmap of proteins differentially expressed in all MPNs compared to controls.

The proteome of *in situ* MPN megakaryocytes was altered compared to controls with 1496 differentially expressed proteins (FDR≤0.05; Figure 2B; Supplemental Table 2). The growth factors FGF2 and EGFL6 were exclusively identified in MPN samples. A subset of proteins were consistently differentially expressed across all MPN subtypes (Figure 2C): upregulated proteins included cell signaling proteins YES1, SOS1, PIK3R6, CD109, cytokines CCL5 and TIMP1, lipid transport and fatty acid biosynthesis proteins such as CD36, LDLR, FADS2, HMGCS1 and ELOVL7, and regulators of cytoskeletal dynamics such as LIMA1, DIAPH1, MACF1, MAP1B and MAP1A; while coagulation factor F5 and immune related proteins including VSIG4, DOCK2, and CD38 were consistently downregulated in megakaryocytes across all MPN subtypes when compared to control megakaryocytes.

#### PMF megakaryocytes are most proteomically divergent to control megakaryocytes

Megakaryocyte atypia is one of the most striking features of PMF.^19^ In keeping with histological observations, PMF patients had the most proteomically distinct megakaryocytes (compared to controls), with 1841 significant (FDR≤0.05) differentially expressed proteins (Figure 3A; Supplemental Table 3). Compared with other MPN and control groups, they exclusively upregulated extracellular matrix (ECM) proteins, including MMP1, THBS2, AGRN, CSPG4, and MXRA5, as well as the actin-regulatory proteins SCIN and CSRP2 and the p53 target CDKN1A.

**Figure 3.**
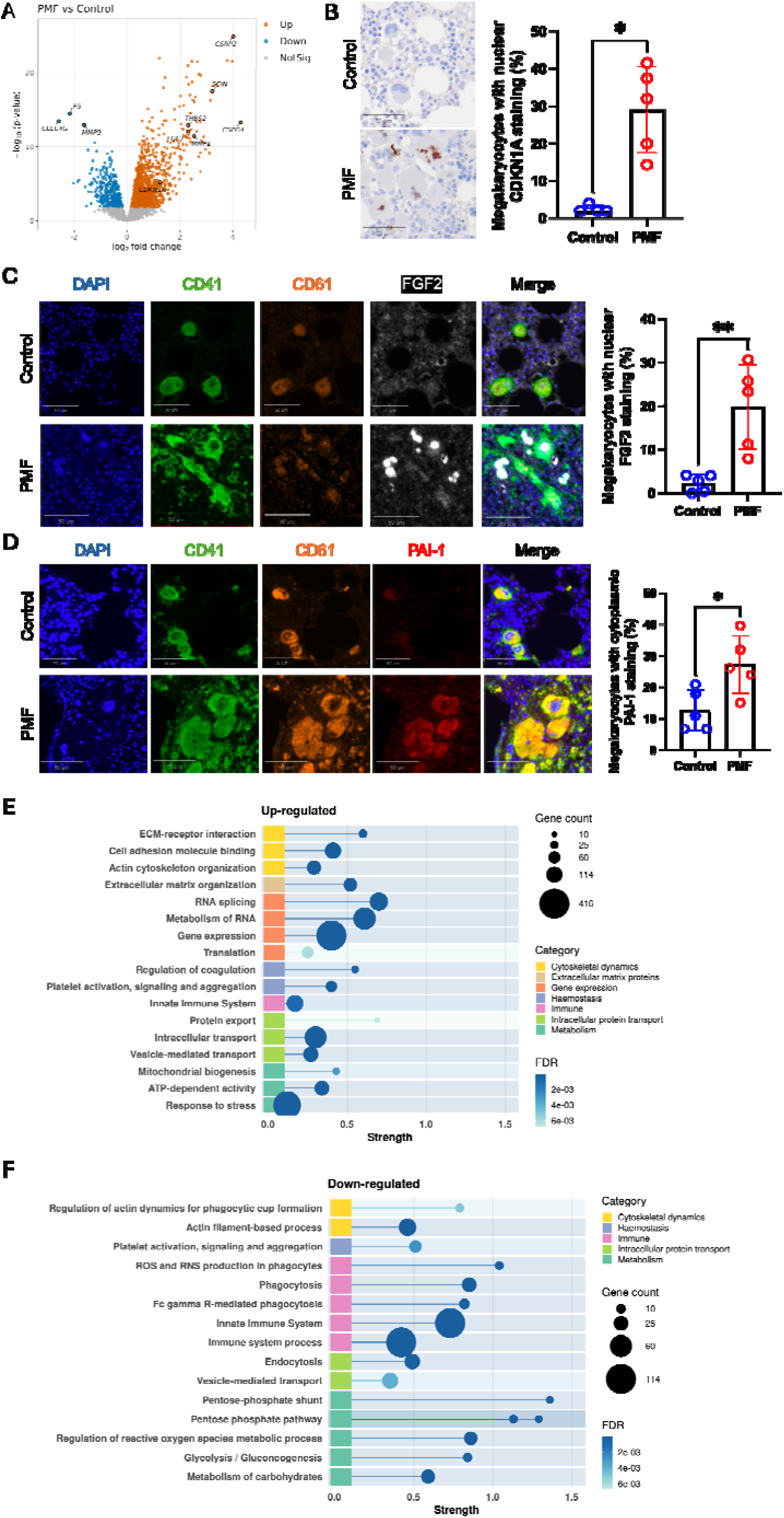
PMF megakaryocytes show substantial proteomic aberrancy. (A) Volcano plot visualizing protein expression in PMF and control megakaryocytes. Orange = increased in PMF; blue = decreased in PMF; grey = not significantly different. (B) IHC validation comparing nuclear CDKN1A expression of megakaryocytes from PMF (n=5) and control (n=4). (C-D) Multiplexed immunofluorescence validation of FGF2 nuclear expression (C) and PAI1 cytoplasmic expression (D) of megakaryocytes from PMF (n=5) and control (n=5). (E-F) Enrichment analysis pathways of upregulated (E) or down regulated (F) proteins in PMF megakaryocytes compared to control megakaryocytes. (B-D) Data represents mean +/- standard deviation.

To confirm our proteomic findings, we validated the expression of three proteins that were differentially expressed in PMF megakaryocytes (CDKN1A, PAI-1 and FGF2) through immunohistochemistry and multiplexed immunofluorescence. We confirmed the upregulation of these proteins in PMF megakaryocytes (Figure 3B-D).

Enrichment analysis was applied to differentially expressed proteins using Gene Ontology^15^, KEGG^16^ and Reactome^17^ databases, and dysregulated pathways were identified and grouped by theme. The most statistically upregulated pathways were related to mRNA splicing and gene expression (Figure 3D). PMF megakaryocytes showed dysregulated cellular metabolism with a shift away from glycolysis towards oxidative phosphorylation, as evidenced by upregulated ATP dependent and mitochondrial biogenesis proteins, and downregulated glycolytic enzymes (Figure 3D-E).

There were increased ECM proteins and cell membrane integrins and non-integrins, suggestive of increased cell membrane-ECM interactions, and corroborated by upregulated cell adhesion terms in pathway analysis (Figure 3E). In keeping with this, cytoskeletal proteins that support stronger adhesion to ECM (CTNNA1, ITGB5, PLEC, EPB41L1/L2, SORBS2/3), and myosin motors (MYH10, MYH11, MYO1B/1D/6) that support actomyosin contractility were increased in PMF megakaryocytes. Conversely, proteins that normally drive lamellipodia, filopodia, and cortical actin dynamics (CEACAM1, CORO1A, CFL1, PFN1, MICAL1, ARPC1B) were downregulated in PMF megakaryocytes, suggesting reduced migratory actin dynamics, and consistent with downregulated innate immune pathways related to phagocytosis. On the other hand, complement proteins (C1QB, C1QC, C2, C4B, C8G, C9, CFB, CFI) and interferon stimulated proteins (IFIT1, ISG15, MX1, MX2) were increased in PMF megakaryocytes, thereby reflecting a state of overall immune dysregulation.

Hemostatic dysfunction was predominantly pro-thrombotic in direction with increased abundance of platelet activation, signaling and aggregation proteins including THBS1, LAT, TBXA2R, PRKAR1B/2B, YES1, RAB27B, SYTL4, ITPR1, ATP2A2, coagulation related proteins such as F13A1and HPSE, and inhibitors of fibrinolysis such as PAI-1 and TIMP1.

### PV and ET megakaryocytes exhibit distinct proteomic signatures when compared to control megakaryocytes

Compared with the marked morphologic atypia of megakaryocytes in PMF, those in PV and ET exhibit more subtle differences, predominantly related to cell size and distribution. Despite these modest morphologic changes, we identified 892 and 880 significant, differentially expressed proteins (FDR ≤ 0.05) in PV and ET megakaryocytes, respectively, compared to control megakaryocytes. (Figure 4A and Figure 4D; Supplemental Tables 4 and 5). Enrichment analysis of differentially expressed proteins revealed dysregulation in immune and hemostatic pathways in both PV and ET megakaryocytes (Figures 4B-C, 4E-F). There was a strong signal for downregulated neutrophil degranulation proteins such as those involved in phagocytosis (PTX3, FGR, SIRPB1, DOCK2) and reactive oxygen species/reactive nitrogen species (ROS/RNS) pathway (MPO, CYBA, NCF1, NCF2, CYBB, NCF4) in ET, while reduced complement proteins (C5, C3, CFP, C6, SERPING1, VSIG4, CD59) were seen in PV megakaryocytes

**Figure 4.**
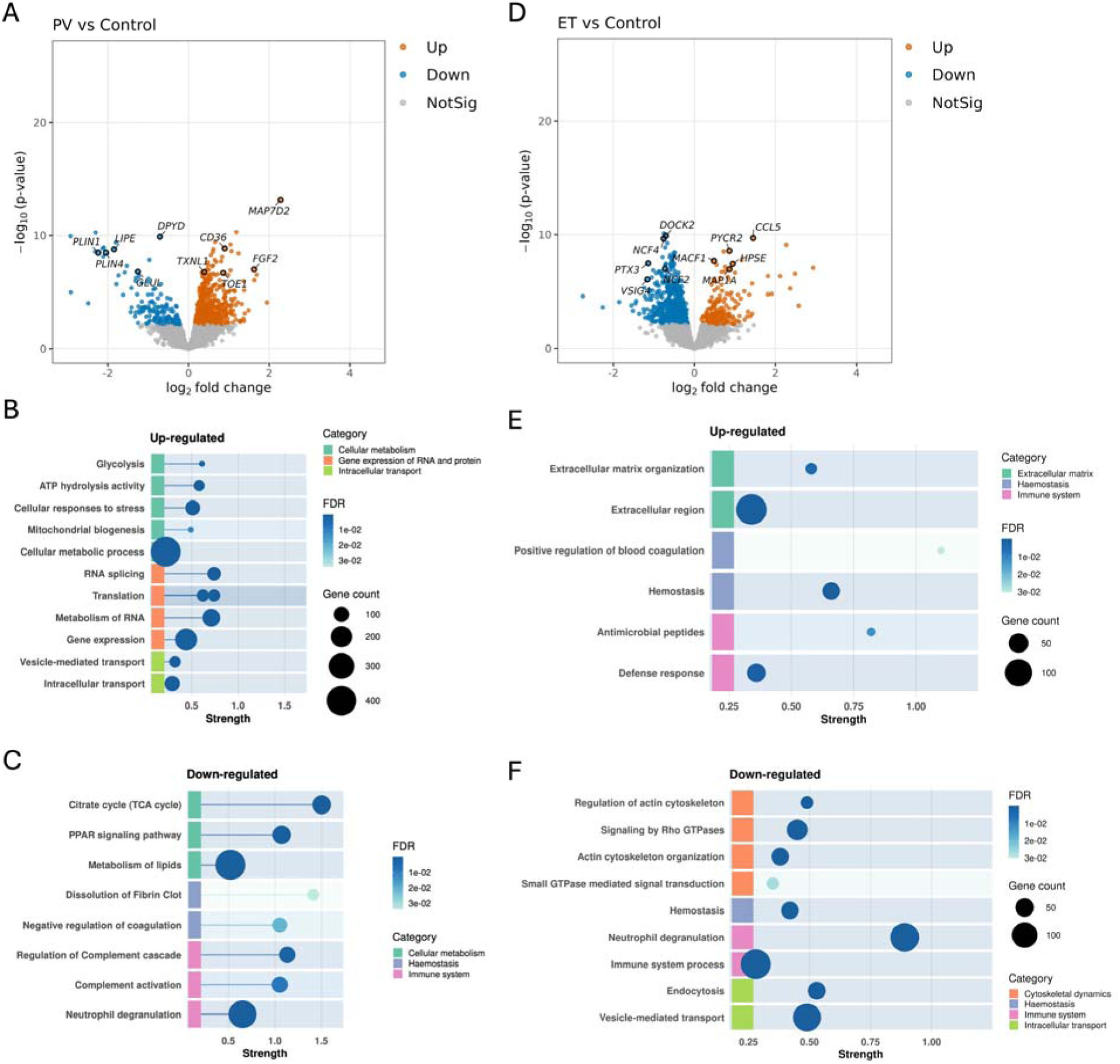
PV and ET megakaryocytes have distinct proteomic signatures. (A) Volcano plot visualizing protein expression in PV and control megakaryocytes. Orange = increased in PV; blue = decreased in PV; grey = not significantly different. (B-C) Enrichment analysis pathways of upregulated (B) or down regulated (C) proteins in PV megakaryocytes compared to control megakaryocytes. (D) Volcano plot visualizing protein expression in ET and control megakaryocytes. Orange = increased in ET; blue = decreased in ET; grey = not significantly different. (E-F) Enrichment analysis pathways of upregulated (E) or down regulated (F) proteins in ET megakaryocytes compared to control megakaryocytes.

With regard to hemostatic pathways, there was increased abundance of platelet activation, signaling and aggregation related proteins in ET, and reduced fibrinolytic proteins such as PLG, SERPINF2, S100A10 in PV, potentially contributing to the prothrombotic state in both diseases (Figure 4B-C, 4E-F). Other downregulated proteins in the hemostasis pathways were predominantly related to immune functions of platelets.

Like PMF megakaryocytes, the most significantly dysregulated pathways in PV megakaryocytes were related to gene expression and protein synthesis, with a corresponding change in intracellular transport and cellular metabolism pathways including increased glycolytic, reduced citric acid cycle and reduced lipid metabolism proteins (Figure 4B-C).

Likewise, ET and PMF megakaryocytes shared some common dysregulated pathways. Both had increased matrix modifying and profibrotic proteins, and reduced cytoskeleton regulatory proteins that are typically responsible for immune cell migration (Figure 4E-F).

### Megakaryocyte proteomic signatures differ between MPN subgroups

Next, we compared the megakaryocyte proteome between the three classical MPNs to define the functional proteomic characteristics underlying morphologic differences of megakaryocytes across the three diseases. We observed significant changes in proteomic landscape between the MPNs with 1385, 1646, and 861 significant (FDR≤0.05) differentially expressed proteins between ET and PV, PMF and ET, and PMF and PV respectively (Figure 5 A-C). We performed sPLS-DA of normalized data to identify proteins that discriminate between the MPN subgroups. sPLS-DA identifies components that are each the linear combination of a relatively small number of protein levels, enabling interpretation of sets of correlated proteins that explain differences between MPN types. Although six components were identified as optimal to separate between MPN subtypes, we found excellent discrimination between MPN subgroups utilizing just two components (Figure 5 D), containing 9 and 90 proteins, respectively (CV AUC = 0.893, 0.937, and 0.968 for ET vs other, PV vs other, and PMF vs other, respectively). Component 1 (9 proteins, found in ECM pathways) separated PMF from ET and PV megakaryocytes, and component 2 (90 proteins, found in gene expression, cellular stress and lipid metabolism pathways) separated ET and PV megakaryocytes (Supplemental Tables 6-7).

**Figure 5.**
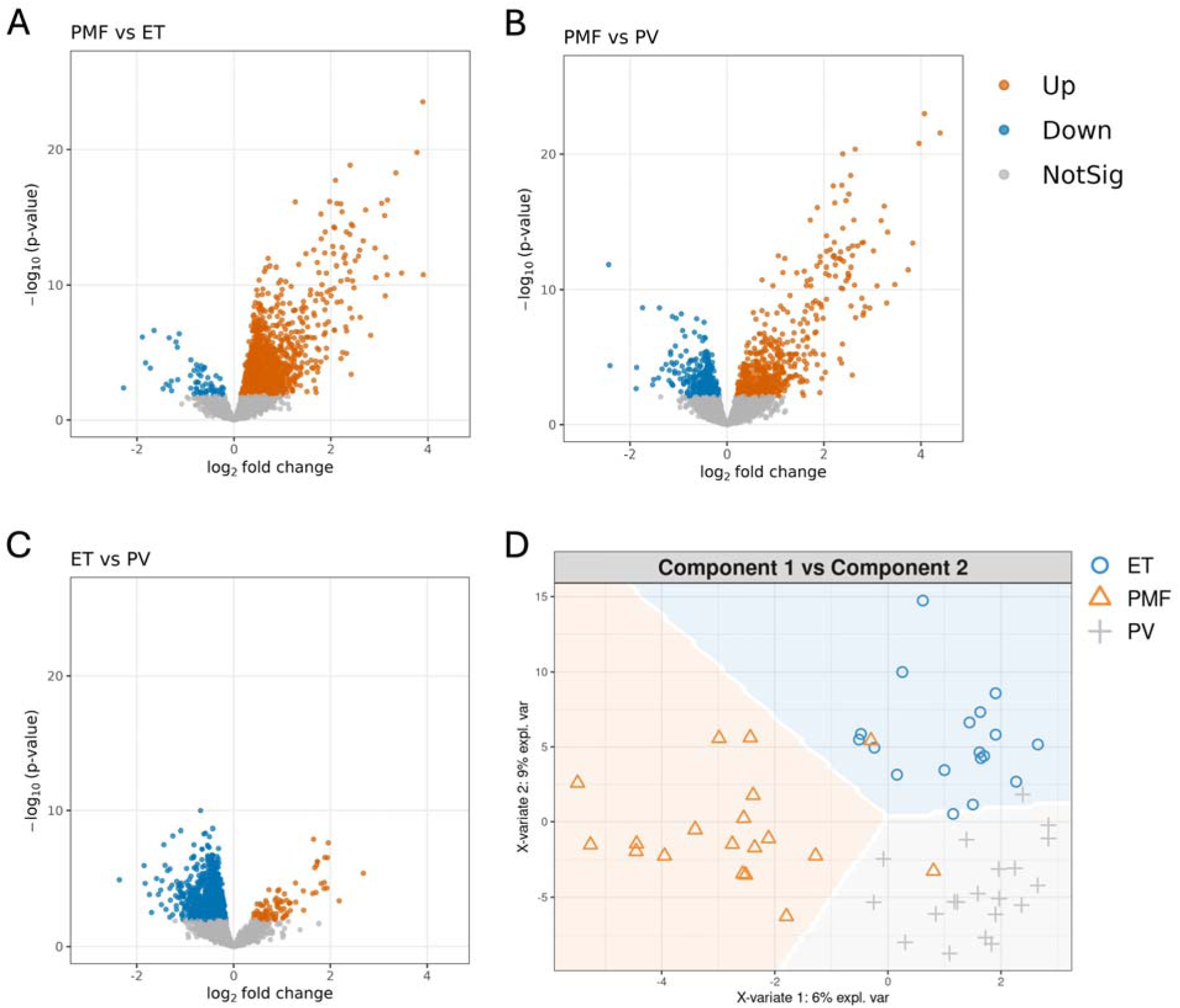
- There are significant proteomic differences between megakaryocytes of the classical MPN. (A-C) Volcano plot of differentially expressed proteins between megakaryocytes of (A) PMF and ET, (B) PMF and PV, and (C) ET and PV. Orange = increased expression; blue = decreased expression; grey = not significantly different. (D) Prediction background plots using component 1 and component 2 derived from sPLS-DA.

## Discussion

Clonal megakaryocytes are drivers of fibrosis and inflammation in the Philadelphia-negative MPNs^19,20^, yet to date comprehensive proteomic studies of bone marrow resident megakaryocytes are lacking. We developed a novel workflow to characterize the proteome of megakaryocytes isolated from archival human bone marrow trephine biopsies of patients and applied this to classical Philadelphia-negative MPN trephines. We show that the proteome of MPN megakaryocytes is significantly aberrant to that of control megakaryocytes, with the most significant changes seen in those obtained from patients with PMF, and that the megakaryocytes across all three conditions harbour distinct proteomic signatures.

Historically, functional proteomic studies of megakaryocytes in health and disease have relied on cells generated *in vitro* from mobilized hematopoietic stem cells (HSCs),^10,21^ reflecting the challenges associated with isolating sufficient numbers of native megakaryocytes for analysis. By integrating deep learning-based image analysis, high-precision laser microdissection, and state-of-the-art mass spectrometry, we substantially improved analytical throughput and sensitivity. This approach enables robust and reproducible proteomic profiling of *in situ* megakaryocytes, capturing cells within their native microenvironment and thereby facilitating the study of disease-relevant phenotypes shaped by a dysregulated niche. We show an enrichment of megakaryocyte specific proteins when compared to matched whole bone marrow trephine proteomic analysis, thereby validating our methodology, which may subsequently be used to profile clinical bone marrow samples in other haematological diseases.

Our findings identify distinct megakaryocyte proteomic signatures across the three MPNs, providing a molecular basis for their well-described morphological differences and potentially aiding disease classification. Across all MPN subtypes, we observed altered expression of proteins involved in immune responses, haemostasis, cytoskeletal organisation, and metabolism. PMF megakaryocytes were distinguished by marked enrichment of structural ECM proteins, including collagens, fibronectin, and thrombospondins. Although expression of ECM components by human megakaryocytes has been established previously^22–24^, evidence for their increased expression in PMF has been limited to a study of *in vitro*-differentiated megakaryocytes from only two patients^24^. Our findings validate and extend these observations in a substantially larger PMF cohort and suggests an active contribution of megakaryocytes to fibrosis in PMF.

Our dataset identified an increase in the expression of the senescence-associated proteins p21 (CDKN1A) and PAI-1 (SERPINE1) in PMF megakaryocytes, which we validated by immunohistochemistry and multiplexed immunofluorescence, respectively. PMF megakaryocytes also exhibited increased expression of several components of the senescence-associated secretory phenotype (SASP)^25^, including TGF-β1, FGF2, TIMP1, MMP1, CCL5, THBS1, FN1, and IGFBP7. In addition, BCL-XL an anti-apoptotic protein associated with the senescent cell phenotype^26^, was increased in PMF megakaryocytes. While p21-mediated senescence is classically induced downstream of TP53,^27,28^ it can also be induced through TGF-β signalling^29^. This pathway may be particularly relevant in PMF, given the established role of TGF-β in the development of myelofibrosis.^30,31^ Furthermore, p21 has been implicated in fibrotic disease through its effects on ECM regulation in pulmonary^32^, and renal fibrosis^33^. Together, these findings raise the possibility that megakaryocyte senescence contributes to the pathological microenvironment of PMF. Determining whether senescent megakaryocytes directly promote fibrosis and/or malignant disease progression, defining the mechanisms underlying this phenotype, and determining if p21 could serve as a therapeutic target, will be an important direction for future studies.

We observed increased expression of interferon-stimulated proteins in PMF megakaryocytes. Treatment data were available for all but one patient with PMF and indicated that none were receiving therapeutic interferon at the time of bone marrow biopsy, suggesting that the observed interferon-associated signature was not attributable to exogenous interferon therapy. Increased interferon signalling has been reported in MPNs in several previous studies.^34–36^ Pegylated interferon-α is an established therapy for MPNs and has disease-modifying activity, in part through targeting of the malignant clone.^37^ It has been proposed that exogenous interferon-α exploits the increased sensitivity of clonal cells to interferon signalling, and promotes the transition of quiescent clonal HSCs into active cycling, differentiation, and apoptosis.^35^ Our data confirms that megakaryocytes in myeloproliferative neoplasms retain evidence of increased interferon signaling, and therefore they are likely to be sensitive to effects of exogenous interferon therapy.

Our findings also identified altered expression of complement components in PMF and PV megakaryocytes. Differential expression of complement proteins have previously been reported in proteomic studies of MPN plasma,^38,39^ and transcriptomic studies of peripheral blood mononuclear cells,^31,37^ both demonstrating increased expression of complement activators and reduced expression of complement inhibitors. In one study, complement proteins were more extensively dysregulated in patients with MPN-associated splanchnic vein thrombosis than in those without thrombosis, raising the possibility of a relationship between complement activation and thrombotic risk in MPNs.^36^ Complement and coagulation pathways are closely interconnected, with bidirectional cross-talk between the two systems, and disorders characterized by dysregulated complement activation, including atypical hemolytic uremic syndrome and paroxysmal nocturnal hemoglobinuria, are associated with a marked thrombotic tendency.^40^ We hypothesize that the altered complement signature observed in MPN megakaryocytes contributes to the prothrombotic phenotype of MPNs.

In summary, we have developed a novel method capable of producing deep quantitative proteomes of *in situ* human megakaryocytes which we applied to MPN patients. Our results strongly suggest dysregulation of homeostatic, immune, and hemostatic functions of MPN megakaryocytes. Our dataset has provided novel disease pathways, biomarkers, and therapeutic targets in MPNs that may be explored further.

## Acknowledgements

The authors thank their patient consumers living with MPNs, Nathalie Cook, Anna Steiner and Jenny Myers for their input and feedback on this work. IR was supported by the Gailey-Lazarus Snowdome Fellowship funded by Epworth Medical Foundation and Snowdome Foundation (SF0033). This work was supported by the National Health and Medical Research Council (GNT2033705 to CB; GNT2009047 to SRP). The contents of this work are the responsibility of the authors and do not reflect the views of the NHMRC. This work was also supported by the Victoria State Government’s Operational Infrastructure Support Program and the NHMRC Independent Research Institute Infrastructure Support Scheme (IRIISS).

## Authorship contributions

I.R., C.B, C.K.Y. and S.-R.P. conceptualized the study; I.R., C.B., E.T., R.A., L.D., S.E.C., P.R., and S.-R.P. designed the research; I.R., S.J., A.P., Y.Z., S.E.C. and S.S. performed the research; I.R., J.Y. and A.M. and analyzed the data; and I.R., C.K.Y, A.P., E.T., R.A., L.D., S.E.C., P.R., J.Y., A.M., S.-R.P and C.B. wrote the manuscript.

