## Supplemental data for "Megakaryocytic proteomic signatures define myeloproliferative neoplasms"

Methods

***Deep learning-assisted*** ***Laser capture microdissection (LCMD)***

Polyethylene naphthalate (PEN) membrane glass slides (4 µm; Leica, 11600288) were pretreated with UV radiation followed by Vectabond® (Vector Laboratories, SP-1800-7). FFPE bone marrow trephine biopsies were sectioned at 4 µm and mounted onto PEN membrane slides.

CD61 immunohistochemistry was performed using a Dako Omnis autostainer. Heat-induced epitope retrieval (HIER) was carried out using high-pH retrieval buffer (EnVision™ FLEX Target Retrieval Solution, High pH; Agilent) at 97 °C for 15 minutes to preserve PEN membrane integrity. Sections were incubated with primary antibody against CD61 (platelet glycoprotein IIIa; mouse monoclonal clone Y2/51, M0753, Dako) at a dilution of 1:50 for 1 hour. Endogenous peroxidase activity was blocked using Dako REAL™ Peroxidase-Blocking Reagent (Agilent, S202386-2) for 4 minutes onboard.

Detection was performed using MACH 4 Mouse Probe (MRH534L/UP534L; Biocare Medical) with a 15-minute onboard incubation, followed by MACH 4 HRP Polymer (RTU; Biocare Medical) for 15 minutes. Visualization was achieved using 3,3′-diaminobenzidine (DAB; Agilent Technologies, GV82511-2) with onboard mixing and incubation for 10 minutes. Sections were counterstained with in-house prepared Mayer’s hematoxylin for 1 minute, followed by bluing reagent (Leica, 3802915) for 1 minute. Slides were dehydrated, cleared, and air-dried.

High resolution images (20x) were obtained with the Navigator module of Leica LAS X (version 3.7.4 Leica Microsystems, Wetzlar, Germany) and tile scans were merged. Deep learning (DL)-based image analysis was carried out using the pixel classifier module of Aivia (Leica Microsystems, Wetzlar, Germany) software to perform automated segmentation, identification and outlining of shapes of CD61 stained megakaryocytes. Using annotated landmarks on the image for spatial orientation, these shapes were transferred to Leica LMD7 software (version 8.3)

Shapes transferred from Aivia (Leica Microsystems, Wetzlar, Germany) to Leica LMD7 (version 8.3) were accurately superimposed upon CD61 stained megakaryocytes from the slide by identifying previously annotated landmarks. Between 200 and 230 megakaryocytes were micro-dissected from each sample into 96 well Eppendorf plates that were pre-filled with 30 µL of acetonitrile (ACN, Fisher Scientific, A9554)) and subsequently stored at negative 20 ^o^C.

***Mass-spectrometry based proteomics***

Samples were re-treated with acetonitrile (ACN, Fisher Scientific, A9554) to aid denaturation and subsequently lyophilized to complete dryness. Dried samples were resuspended in 10 µL of 8 M urea (Novachem, 216-00185) in 100 mM Tris pH 7.5 (Urea-Tris Buffer) containing 10 mM DTT (Sigma, D5545) and reduced at 37 °C for 45 min. Alkylation was performed by adding 5 µL of Urea-Tris buffer containing 60 mM iodoacetamide (IAM, Sigma, I1149), followed by incubation at room temperature in the dark for 45 min. Excess IAM was quenched with 5 µL of the DTT-containing Urea-Tris buffer. For enzymatic digestion, samples were diluted with 20 µl of 100 mM Tris to reduce urea concentration to >4 M prior to the addition of 10ul of diluted Lys-C (2.5 ng/µl, Wako, 129–02541) in 100 mM Tris, pH 7.5 at 37 °C for 2 h. Subsequently 40 µl of 100 mM Tris was added to reduce urea concentration to >2 M for trypsin digestion. 10µl of diluted trypsin (50 ng/ul, SOLu‐Trypsin, EMS0004) was added and incubated overnight at 37 °C. The following day, peptide digests were acidified with 10 µL of 20% formic acid and purified using AffiniSep C18 stage tips (PM Separations, Tips-C18.T3.200.96 AFF). Tips were conditioned with acetonitrile, 50% ACN/0.1% formic acid (FA), and 0.1% FA before sample loading, followed by washing twice with 0.1% FA and elution with 50% ACN/0.1% FA. Peptides were lyophilized to dryness using a CentriVap (Labconco) before being reconstituted in 4 µL 0.1% FA/2% ACN ready for mass spectrometry analysis. Reconstituted peptides (3 µL, approximately 100 ng) were separated using reverse-phase liquid chromatography on a 15 cm C18 fused silica column with an integrated emitter tip (IonOpticks, ID 75 µm, OD 360 µm, 1.6 µm C18 beads) on a Neo Vanquish HPLC system coupled to a Orbitrap Astral mass spectrometer (Thermo Fisher Scientific). Peptides were analyzed on a 25 min linear analytical gradient of increasing buffer B (80% ACN, 0.1% FA) from 2 to 34%. Data was acquired in a data independent acquisition (DIA) mode. The MS1 settings were as follows: Orbitrap resolution 240,000; scan range m/z 380-980; AGC target 500%. The DIA parameters were as follows: isolation window: custom; HCD collision energy: 25%; scan range: m/z 145-1450; maximum injection time: 3 ms; AGC target: 800%. The mass spectrometry proteomics data have been deposited to the ProteomeXchange Consortium via the PRIDE partner repository with the dataset identifier PXD073839.

***Immunohistochemistry***

IHC staining was performed with primary antibody against CDKN1a (p21; rabbit monoclonal clone 12D1, 2947, CST) on the Dako Omnis (Agilent) automated staining platform. Formalin fixed, paraffin embedded tissue sections were deparaffinized, rehydrated, and subjected to heat-induced epitope retrieval using EDTA buffer (EnVision FLEX TRS High pH, Cat# GV80411-2) at 97 ^o^C for 30 mins.

Endogenous peroxidase activity was blocked using Dako REAL Peroxidase-Blocking Reagent (Cat# S202386-2) for 4 minutes. Tissue sections were then incubated for 1 hour with rabbit anti-p21 antibody (Cell Signaling Technology, Clone 12D1, Cat# 2947, dilution 1:50). Detection was performed using a ready to use anti-rabbit horseradish peroxidase (HRP) conjugated polymer detection system (Agilent, Cat# K400311-2). Slides were subsequently incubated with DAB chromogen for 10 minutes.

Following completion of the staining run, slides were removed from the Dako Omnis and transferred to a Leica Autostainer XL for counterstaining. Sections were counterstained with WEHI in-house prepared Mayer’s haematoxylin for 1 minute, followed by Bluing Reagent (Leica, Cat# 3802915) for 1 minute. Slides were then dehydrated and coverslipped using a Leica CV5030 coverslipper.

***Multiplexed immunofluorescence***

OPAL multiplex staining was performed using a BOND RX Fully Automated Research Stainer (Leica Biosystems). Following onboard deparaffinization and rehydration, antigen retrieval was carried out using either citrate-based Bond Epitope Retrieval Solution 1 (ER1; Leica Biosystems, cat. no. AR9661) or EDTA-based Bond Epitope Retrieval Solution 2 (ER2; Leica Biosystems, cat. no. AR9640), depending on the antibody. Tissue sections were blocked with Background Sniper protein blocking solution (Biocare Medical, SKU BS966) for 10 min, followed by endogenous peroxidase blocking using Dako REAL Peroxidase-Blocking Reagent (Agilent, cat. no. S2023) for 10 min.

For the multiplex panels, tissue sections were incubated sequentially with primary antibodies (anti-CD61 [Dako M0753]; anti-CD41 [Abcam ab134131]; anti-PAI-1 [Abcam ab318286]; anti-FGF2 [Abcam ab92337]), followed by detection using anti-rabbit or anti-mouse horseradish peroxidase (HRP) polymer and visualization with Opal fluorophores (Akoya Biosciences). Between staining cycles, antibody stripping was performed using antigen retrieval solutions, with retrieval conditions optimized for each antibody.

After completion of all staining cycles, tissue sections were counterstained with DAPI (Akoya Biosciences, cat. no. FP1490; 3 drops/mL) for 5 min. Slides were then removed from the autostainer and treated with TrueBlack Lipofuscin Autofluorescence Quencher (Biotium, cat. no. 23007) for 30 s before mounting with ProLong Diamond Antifade Mountant (Invitrogen, cat. no. P36961). Whole slide images were acquired using a PhenoImager HT scanner (Akoya Biosciences)
